# Microliter spotting and micro-colony observation: a rapid and simple approach for counting bacterial colony forming units

**DOI:** 10.1101/2022.01.26.477842

**Authors:** Shuvam Bhuyan, Mohit Yadav, Shubhra Jyoti Giri, Shuhada Begum, Saurav Das, Akash Phukan, Pratiksha Priyadarshani, Sharmilee Sarkar, Anurag Jayswal, Kristi Kabyashree, Aditya Kumar, Manabendra Mandal, Suvendra Kumar Ray

## Abstract

For enumerating viable bacteria, traditional dilution plating to count colony forming units (CFU) has always been the preferred method in microbiology owing to its simplicity, albeit laborious and time-consuming. Similar CFU counts can be obtained by quantifying growing micro-colonies in conjunction with the perks of a microscope. Here, we employed a simple method of five microliter spotting of differently diluted bacterial culture multiple times on a single Petri dish followed by finding out CFU by counting micro-colonies using a phase-contrast microscope. In this method, the CFU of an *Escherichia coli* culture can be estimated within a four-hour period. Further, within a ten-hour period, CFU in a culture of *Ralstonia solanacearum*, a bacterium with a generation time of around 2 h, can be estimated. The CFU number determined by micro-colonies observed is comparable with that obtained by the dilution plating method. Micro-colonies number observed in the early hours of growth (2 h in case of *E. coli* and 8 h in case of *R. solanacearum*) were found to remain consistent at later hours, though there was a noticeable increase in the size of the colonies. It suggested that micro-colonies observed in the early hours indeed represent the bacterial number in the culture. Practical applications to this counting method were employed in studying the rifampicin-resistant mutation rate as well as performing the fluctuation test in *E. coli*. The method described here results in a 90% reduction of labour, time and resources. Thus, the method is likely to be adopted by many microbiologists in their routine laboratory research.

## Introduction

Quantifying bacterial numbers by colony forming units (CFU) by employing serial dilution and spread plate techniques is an age-old fundamental microbiology technique used in research and diagnostics (Jett et al., 1997; McNulty and Dunn, 1999; Yadav and Rathore, 2022). Though popular among microbiologists because of its ease, the method demands large volumes of growth media, time and labour. Besides, working with bacteria having a slow generation time such as *Ralstonia solanacearum*, which takes ~48 h to form a full-grown colony on a solid agar nutrient medium only doubles the misery (Kumar et al., 2013; Xu et al., 1990). Several other methods for enumerating the number of bacteria in culture have come to light over the years. These include the measurement of absorbance using a spectrophotometer or a plate reader; flow cytometry, microscopy in conjunction with staining, qPCR and several microarray-based methods with each method having its own advantages over others (Hsieh et al., 2018; Koch, 2014; Pan et al., 2014; Stevenson et al., 2016). Despite all advancements and fast approaches for enumerating bacteria in a culture (Jett et al., 1997; McNulty and Dunn, 1999; Hsieh et al., 2018), CFU counting has been the most dependable and popular method among microbiologists because of its viable nature and visual impact. Therefore, it will be a big boon for microbiologists to count CFU in a bacterial culture with reduced labour, time and cost.

In this study, we are describing a subtle method of enumerating bacteria in culture by observing micro-colonies under a microscope. The basic principle of this method is to spot a small amount (~ 5 μl) of a serially diluted bacterial culture followed by imaging and quantification of the micro-colonies obtained using the microscope (Ray et al., 2015). In an *E. coli* culture, CFU can be found within four hours after observing the micro-colonies by this method. Similarly, in the case of *R. solanacearum*, a bacterium with a generation time of about 2 h, CFU can be found within ten hours after observing the micro-colonies. The method is simple to use, scalable and cost-effective, which maintains the benefits of a microscopic platform. Further, validation of this method was done by comparison with other standard quantification approaches. It is pertinent to note that the micro-colonies count method is done by spotting a few microliters, which enables us to analyse different dilutions in a single petri dish. Therefore, the number of CFU observed under the microscope can be confirmed by observing the CFU through the naked eye in later days depending on the growth rate of the bacteria.

Owing to its easy and simple approach, enumeration of the total population of *E. coli* in mutant frequency calculation and the determination of the number of *E. coli* generations was aided using this microlitre spotting and micro-colony counting method. The data obtained was subsequently substituted into the mutation rate formula to calculate it. In addition, the classical fluctuation test by Luria and Delbruck, (1943) has been studied by analyzing rifampicin-resistant (Rif^r^) mutants in *E. coli* culture.

## Materials and methods

### Strains and growth conditions

*E. coli* DH5α and BL21 cells were grown in Luria–Bertani (LB) medium at 37°C. *R. solanacearum* F1C1 cells were grown in BG medium (which contains 1% peptone, 0.1% yeast extract and 0.1% casamino acids) supplemented with 5 g/L glucose at 28 °C. For solid medium, agar 1.5% was used.

### Colony-forming unit (CFU) count

The bacterial strains were inoculated in their respective liquid medium and incubated at their optimal growth conditions for 6 h in *E. coli* and overnight in *R. solanacearum* with a 150 rpm shaking condition. For counting CFU, the overnight grown culture of *E. coli* cells was serially diluted and spotted on LB agar plates as illustrated in Fig. 1. First, 100 μL overnight grown culture of *E. coli* was diluted in 900 μL of sterile distilled water. Since the dilution was one part sample to nine parts diluents, it was referred to as a 10^1^ dilution. Further, we took 100 μL of the 10^1^ dilutions and added it to 900 μL of the sterile distilled water. This dilution was a 10^2^ dilution. Lastly, we made a 10^10^ dilution by taking 100 μL of the 10^9^ dilutions and adding 900 μL of the sterile distilled water. For each dilution, we now had 1 mL of diluted bacterial culture. After two hours of incubation at 37 °C, micro-colonies were counted by using Life technologies EVOS FL inverted microscope with 40X magnification using the 4/10 PH condenser annulus. Under the microscope, the area covered by the cells was calculated by extrapolating the length of the 1000 μm scale provided. Thus, we calculated the total CFU in the area covered by a 5 μL spot and subsequently in 1 mL of *E. coli* culture.

**Fig. 1:**
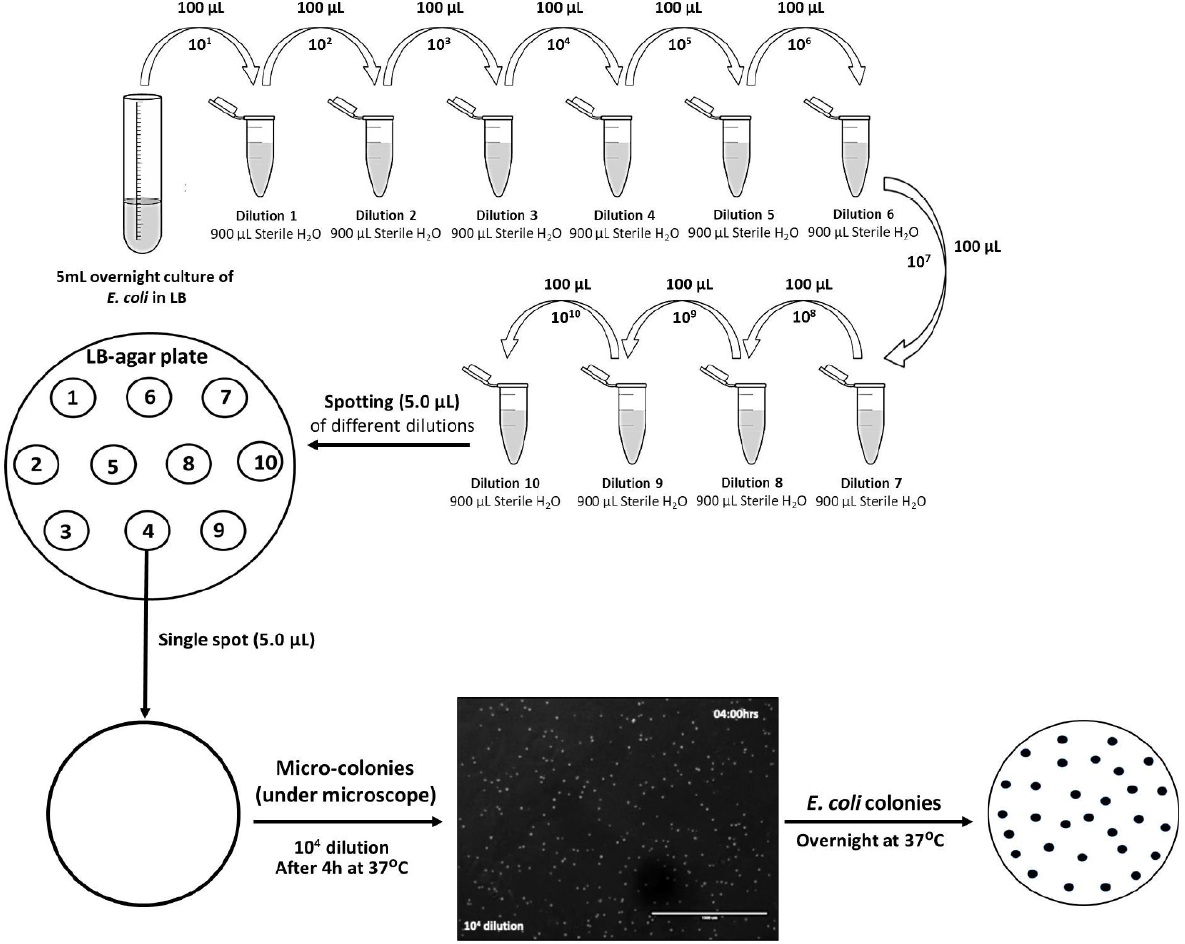
Schematic representation of the microcolonies counting method. In a given bacterial culture considered for counting CFU, dilution of the culture was performed as shown in the figure. 100 μl of the culture was added to 900 μl of water to make it 10-fold (10^1^) dilution, from the diluted culture 100 μl was taken and added to 900 μl of water to dilute it another 10 times. The culture was diluted 10^10^ fold. Only 5 μl from different dilutions were spotted on an LB-Agar plate in the case of *E. coli*. Approximately, ten such spots can be made on Petri plates (90 mm diameter). After 2 h of incubation, micro-colonies were observed at 40x magnification. Three different sections of 10^4^ dilutions spotted have been shown here. Distinct colonies could be observed.

This micro-colony counting was further validated by on-plate CFU counting of overnight incubation at 37°C. The mean value of ten independent spots was used to show the growth parameters. A similar approach was used to count CFU in the case of *R. solanacearum* culture. However, the culture was incubated at 28°C. As *R. solanacearum* is a slow-growing bacterium with a generation time of about 2 h, its micro-colonies at different dilutions could be observed distinctly at 40X magnification in 10 h of incubation after spotting. The bacterial CFU was also counted on Life technologies EVOS FL inverted microscope with 40X magnification using the 4/10 PH condenser annulus.

### CFU from the micro-colony counting method

Micro-colonies observed under the microscope of different dilutions such as 10^4^, 10^5^, 10^6^ -fold etc. were counted at different time intervals manually. In the case of *E. coli* observation of micro-colonies was done from 2 h onwards up to 8 h at every 2 h intervals in the EVOS FL microscope with an objective 40X magnification. The area covered under the microscope was calculated from the scale bar 1000 μm provided in the microscope. The total area covered under the microscope was calculated as 5.67 mm^2^. A 5 μl culture covers an average area of 19.62 mm^2^ (a circle with an average diameter of 5 mm). From a single spot of bacterial culture, a minimum of three sections (each 5.67 mm^2^) were counted and the average number of colonies per section was found, which was then multiplied by 3.46 to find out the number of bacterial colonies in 5 μl of the culture at that dilution. The same approach was used to find CFU at different dilutions used for spotting the culture. In the case of *R. solanacearum*, observation of micro-colonies was done from 8 h onwards up to 14 h at every 2 h intervals.

### CFU from microliter spotting

After observing the micro-colonies under the microscope in their early hours of growth, the Petri dishes were kept back in the incubator. After overnight growth (in the case of *E. coli*) colonies could be visible by naked eyes. In the case of 10^6^ dilutions, an average of 15 colonies per spot can be observed, while in the case of 10^7^ dilutions average of 2 colonies per spot was observed. This method proved that the observations made earlier through micro-colonies counting were correct.

### CFU from the spread plate method

In the usual spread plate method to count CFU, 100 μl of overnight grown serially diluted bacterial cultures were evenly distributed on the solid growth medium by using a glass spreader. The Petri dishes were kept in the incubator and after overnight incubation, we observed the colonies with naked eyes. All experiments were performed in triplicates.

### Calculation of mutation frequency

To find out the mutation frequency, the number of mutant colonies grown on antibiotic-supplemented growth media was counted out of the total number of bacterial populations (CFU), which is given by the number of mutant colonies divided by the total bacterial CFU.

### Calculation of number of generations *(n)*

The number of times that cells divide between the initial and final time is the number of generations (n) for that time interval. This was calculated as-

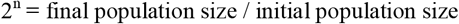

### Generation time calculation

Generation time (t_g_) was calculated by dividing the total time of incubation (T) by the number of generations (n). The formula is given by-

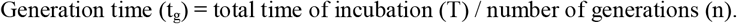

### Mutant rate calculation

The mutation rate was calculated by using the following formula-

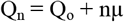

Where ‘Q_n_’ is the final mutation frequency, ‘Q_o_’ is the initial mutation frequency, ‘n’ is the number of generations and ‘μ’ is the mutation rate (Maloy et al., 2008).

### Rifampicin resistance (Rif^r^) mutant isolation

Initially, *E. coli* BL21 was streaked on an LB agar plate from a previously-stored glycerol stock maintained at −80 °C. A single colony from the plate was taken and inoculated in 5 ml of LB broth and kept under incubation at 37 °C for 24 h. Then 100ml of 1% initial culture broth was inoculated and kept under incubation at 37 °C. After 1 h of bacterial growth/incubation, 10 ml of the culture was taken and spun down at 4000 rpm for 15 mins at 20 °C. Decanting the supernatant the pallet was resuspended on 1ml of sterile distilled H_2_O. Out of 1 ml of cell suspension, 100 μl was spread on each plate of LB agar supplemented with 50 μg/ml rifampicin. This was followed by 1 hour of intervals up to 4 hours.

### Fluctuation test

Overnight-grown *E. coli* DH5α was inoculated in 5 ml of LB medium and kept under incubation overnight at 37 □ temperature to get a saturated primary bacterial culture. 100 ml of LB broth was then inoculated with 10 μL of primary culture and it was marked as secondary culture. Soon after inoculation, the secondary culture was distributed in 20 different test tubes, 2 ml in each; the remaining culture was kept on the flask itself and incubated at 37 □. After a continuous incubation of 7 h (~13 generations), cultures were transferred into centrifuge tubes and at the same time, 20 microfuge tubes were filled with cultures taken from the flask. Centrifugation was done at 4000 rpm for 10 min at 20 □. Pellets obtained in each tube were resuspended in 100 μL of sterile distilled water. Around 5 spots were spotted with 10 μL of resuspended pellet on LB plate containing rifampicin antibiotic from each tube. Plates were incubated at 37 □ for 24 h and counted for the rifampicin resistance in mutant colonies on plates [Fig. 5].

## Results and Discussion

We calculated the micro-colonies in an *E. coli* culture as described in the Materials and Methods section. We observed the micro-colonies of *E. coli* by the microscope from ten different spots of different dilutions such as 10^4^, 10^5^, and 10^6^ from 2 h onwards after incubation till 8 h after every 2 h interval [SFig1a-1c]. The number of micro-colonies observed and CFU calculated are listed in Table 1. After 2 h of incubation, the micro-colonies counts were 2.22 x 10^9^ and 3.62 x 10^9^ for the 10^4^ and 10^5^ dilution spots, respectively. Subsequently, in 4 h of incubation period at 37 °C, the micro-colonies counts were calculated as 1.99 x 10^9^, 2.40 x 10^9^ and 4.98 x 10^9^ for the 10^4^, 10^5^ and 10^6^ dilution spots, respectively [Fig. 2a, Table 1]. After 6 h of the incubation period, the micro-colonies counts were found to be 1.8 x 10^9^, 2.35 x 10^9^ and 3.04 x 10^9^ for the 10^4^, 10^5^ and 10^6^ dilution spots, respectively. While, in 8 h of incubation, the micro-colonies counts were calculated as 1.26 x 10^9^, 2.10 x 10^9^ and 3.46 x 10^9^ for the 10^4^, 10^5^ and 10^6^ dilution spots, respectively. Further, these micro-colonies counts were confirmed by spot dilution [Fig 4A] and plate spreading [SFig 3E-3H] methods and results showed that almost similar CFU counts were observed with different dilutions at different time periods as listed in Table 3. It supported that micro-colonies count can represent the CFU in a culture in the case of *E. coli*.

**Table 1:** CFU calculated in *E. coli* culture by the micro-colony counting method.

| Dilution | Time (h) | CFU/mL |
| --- | --- | --- |
| $10^4$ | 2 | $2.22 \times 10^9$ |
| | 4 | $1.99 \times 10^9$ |
| | 6 | $1.80 \times 10^9$ |
| | 8 | $1.26 \times 10^9$ |
| $10^5$ | 2 | $3.62 \times 10^9$ |
| | 4 | $2.40 \times 10^9$ |
| | 6 | $2.35 \times 10^9$ |
| | 8 | $2.10 \times 10^9$ |
| $10^6$ | 4 | $4.98 \times 10^9$ |
| | 6 | $3.04 \times 10^9$ |
| | 8 | $3.46 \times 10^9$ |

**Fig 2.**
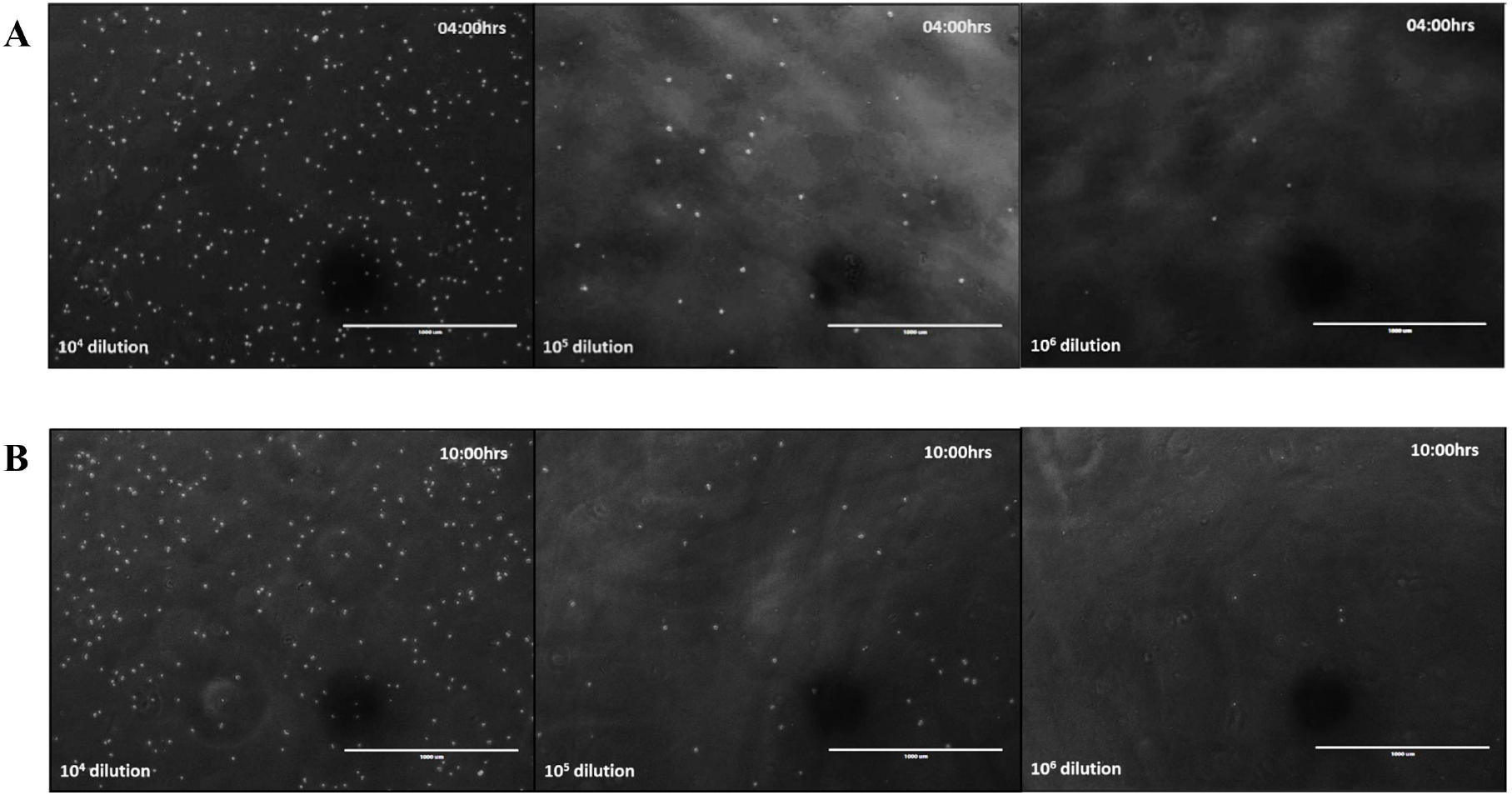
A. *E. coli* microcolonies number observed at different dilutions *E. coli* microcolonies were observed at different dilutions such as 10^4^, 10^5^ and 10^6^ after 4 h incubation. Microcolonies formed are distinctly observed at different dilutions. The number decreases with an increase in dilution. This suggests that CFU count is possible after 4 h of incubation in the case of *E. coli*. B. *R. solanacearum* microcolonies at different dilutions *R. solanacearum* microcolonies were observed at different dilutions such as 10^4^, 10^5^ and 10^6^ after 10 h incubation. Microcolonies formed are distinctly observed at different dilutions. The number decreases with an increase in dilution. This suggests that CFU count is possible after 10 h of incubation in the case of *R. solanacearum*.

**Fig 3.**
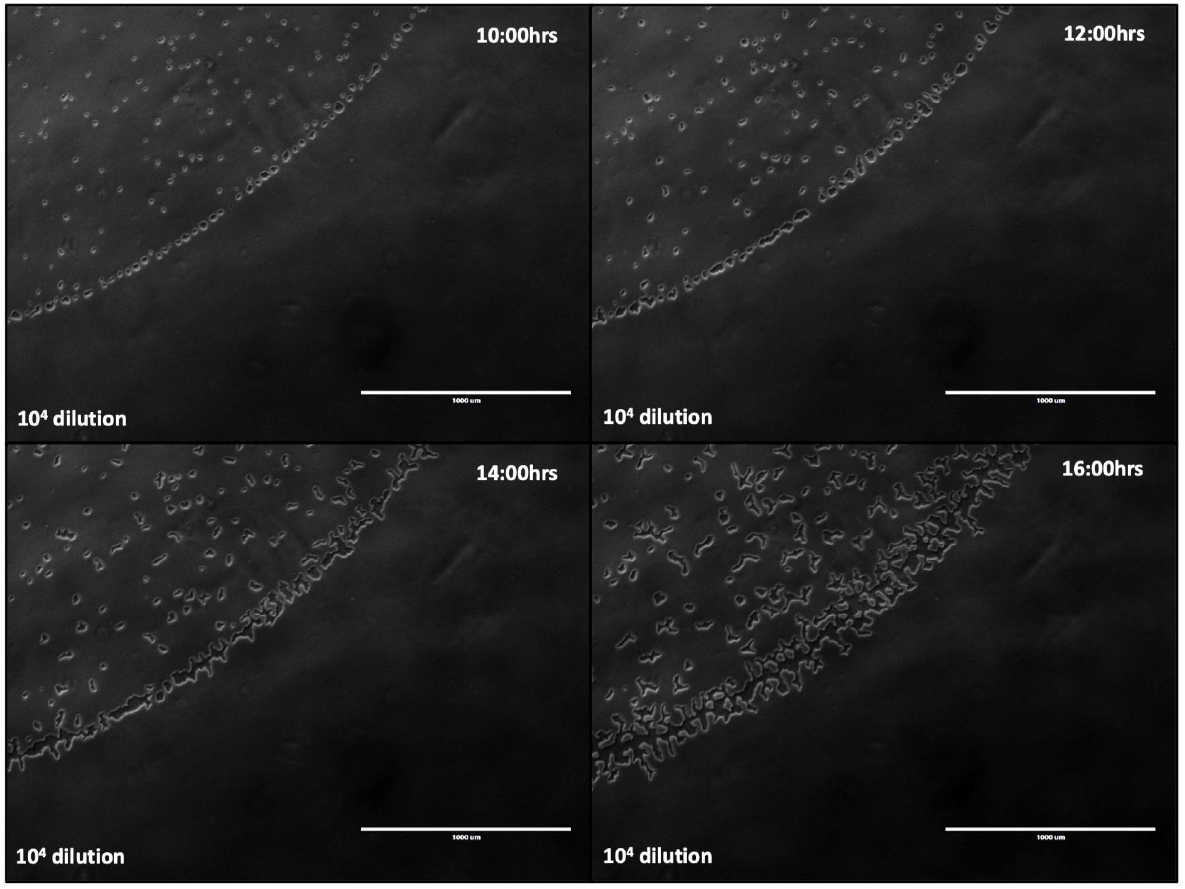
*R. solanacearum* microcolonies number observed at different time points. At 8 h, 10 h, 12 h and 14 h duration microcolonies were observed for 10^4^ dilutions of *R. solanacearum* culture. Counted microcolonies were found to be similar at different time points though there is growth in colony size. Further, the number of CFU estimated by counting the microcolonies at different dilutions is observed to be similar.

**Fig 4.**
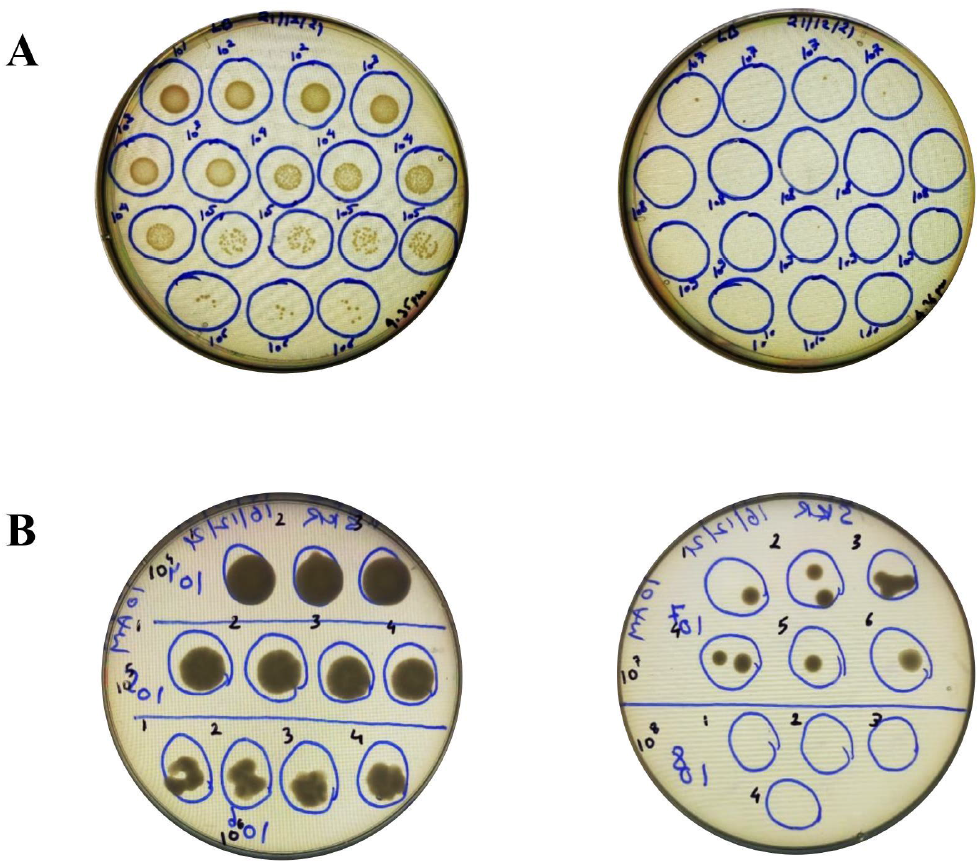
A. *E. coli* colonies at different dilutions by spotting method *E. coli* colonies were observed after 16 h of spotting. At 10^5^ and 10^6^ dilutions, distinct colonies can be observed. B. *R. solanacearum* colonies at different dilutions by spotting method *R. solanacearum* colonies were observed after 48 h of spotting. At 10^7^ dilution, distinct colonies can be observed.

Likewise, the CFU count using microliter spotting and micro-colony counting was done in *R. solanacearum*, a bacterium with a generation time of about 2 h, as well. Micro-colonies could be observed from 8 h onwards at 10^4^ dilution spotting. However, it could be counted properly after 10 h of spotting. Fig 2b demonstrates micro-colonies of *R. solanacearum* after 10 h of spotting of dilutions such as 10^4^, 10^5^, and 10^6^. It is pertinent to note that at this hour of growth it is not easy to find colonies in the case of 10^7^ dilutions as the number of colonies is very low. We observed that the micro-colonies of *R. solanacearum* by the microscope from ten different spots of 10^4^, 10^5^ and 10^6^ each from 8 h of incubation till 14 h after every 2 h interval [SFig 2a-2c]. The number of micro-colonies observed and CFU calculated are listed in Table 2. After 8 h of incubation, the micro-colonies counts were 1.74 x 10^9^, 2.86 x 10^9^ and 3.72 x 10^9^ for the 10^4^, 10^5^ and 10^6^ dilution spots, respectively. Subsequently, in 10 h of incubation period at 28 °C, the micro-colonies counts were calculated as 1.78 x 10^9^, 2.86 x 10^9^ and 3.85 x 10^9^ for the 10^4^, 10^5^ and 10^6^ dilution spots, respectively [Table 2]. After 12 h of the incubation period, the micro-colonies counts were found out 1.76 x 10^9^, 2.79 x 10^9^ and 3.99 x 10^9^ for the 10^4^, 10^5^ and 10^6^ dilution spots respectively. While, in 14 h of incubation, the micro-colonies counts were calculated as 1.51 x 10^9^, 2.57 x 10^9^ and 3.79 x 10^9^ for the 10^4^, 10^5^ and 10^6^ dilution spots, respectively. Similar results were obtained with the plate spreading method [Table 3, SFig 3A-3D]. We further counted 10 spots for 10^6^ and 10^7^ dilutions each, as well as 10 plates each of dilutions 10^6^ and 10^7^ of *E. coli* and the observation revealed that the coefficient of variance was comparable by both the spotting method and the dilution plating method, respectively [Table 4]. The size of micro-colonies has increased at different time periods; however, these results show that there is a similarity in the calculated number of micro-colonies at different time periods [Fig. 3]. It supported that micro-colonies count could represent the CFU in a culture in the case of *R. solanacearum*. In *R. solanacearum*, the micro-colonies grow in size, exhibit twitching motility and aggregate together [Fig. 3]. Twitching motility is, however, absent in *E. coli*.

**Table 2:** CFU calculate in *R. solanacearum* by micro-colony counting method.

| Dilution | Time (h) | CFU/ml |
| --- | --- | --- |
| $10^4$ | 8 | $1.74 \times 10^9$ |
| | 10 | $1.78 \times 10^9$ |
| | 12 | $1.76 \times 10^9$ |
| | 14 | $1.51 \times 10^9$ |
| $10^5$ | 8 | $2.86 \times 10^9$ |
| | 10 | $2.86 \times 10^9$ |
| | 12 | $2.79 \times 10^9$ |
| | 14 | $2.57 \times 10^9$ |
| $10^6$ | 8 | $3.72 \times 10^9$ |
| | 10 | $3.85 \times 10^9$ |
| | 12 | $3.99 \times 10^9$ |
| | 14 | $3.79 \times 10^9$ |

**Table 3:** Comparison of CFU calculation by 5 μl spotting and 100 μl spreading method.

| Dilutions | *R. solanacearum* |  | *E. coli* |  |
| --- | --- | --- | --- | --- |
|  | Dilution spotting | Spread plate | Dilution spotting | Spread plate |
| $10^4$ | $2.1 \times 10^9$ cells | Merged colonies | Merged colonies | |
| $10^5$ | $2.3 \times 10^9$ cells | Merged colonies | $6.04 \times 10^8$ | $4.5 \times 10^8$ |
| $10^6$ | $3.5 \times 10^9$ | $10^9$ | $6.2 \times 10^8$ | $7.1 \times 10^8$ |
| $10^7$ | $7 \times 10^9$ | $1.1 \times 10^9$ | $1.6 \times 10^9$ | $7 \times 10^8$ |
| $10^8$ | No colonies found | $6.7 \times 10^8$ | $4 \times 10^9$ | $6.7 \times 10^8$ |

**Table 4:** Comparison of visible colonies observed by both the methods in *E. coli*.

|  | 10<sup>6</sup> dilution | 10<sup>7</sup> dilution |  | 10<sup>6</sup> dilution | 10<sup>7</sup> dilution |
| --- | --- | --- | --- | --- | --- |
| S | 5μL | 5μL | S | 100μL | 100μL |
| P | 20 | 2 | P | 121 | 13 |
| O | 21 | 1 | R | 233 | 13 |
| T | 11 | 0 | E | 106 | 30 |
| T | 15 | 1 | A | 102 | 13 |
| I | 19 | 1 | D | 156 | 14 |
| N | 16 | 0 | I | 80 | 19 |
| G | 9 | 5 | N | 199 | 21 |
|  | 14 | 1 | G | 242 | 29 |
|  | 11 | 3 |  | 210 | 23 |
|  | 17 | 1 |  | 205 | 27 |
| MEAN | 15.3 | 1.5 | MEAN | 165.4 | 20.2 |
| SD | 4.08 | 1.51 | SD | 59.65 | 6.86 |
| CV | 0.27 | 1.01 | CV | 0.36 | 0.34 |

Serial dilution is an easy and convenient approach for measuring the viable bacterial concentration in a culture. In the early days of microbiology, enumerating bacterial numbers in culture was made in test tubes by finding out the minimum dilution at which no bacterial growth was observed (Koch, 1883). Usually, turbidity development in the medium used to be measured as bacterial growth [SFig 4]. Later with the use of agar, colony formation by bacteria could be observed in solid agar medium (Breed and Dotterrer, 1916). With the availability of micropipettes, which can pipette out liquids of volume 1.0 or 2.0 μl with accuracy, it became an aid for us to think of counting bacteria in 5 μl volume. The findings that micro-colonies number remain similar at various points of time at a specific dilution supported that the method can be used to count bacteria in culture by observing micro-colonies. In contrast to the conventional spread plate method, calculating CFU by dilution spotting is both time-saving and economical. While we could calculate *R. solanacearum* and *E. coli* CFU using dilution spotting at 10 h and 4 h respectively, we could obtain similar results for *R. solanacearum* and *E. coli* using the spread plate method at only 48 h and 16 h of incubation respectively.

Dilution spotting and subsequent back calculations are a fast and easy technique to enumerate the bacterial population in culture. Besides, CFU can be estimated from different dilutions, such as 10^4^, 10^5^, 10^6^ and 10^7^. Counting CFU from 10^4^ and 10^5^ dilution by spread plate method is difficult because the colonies merged with each other due to their high number. Also, less variation of CFU numbers across different spots of the same dilution is observed. A dilution spot of 10^5^ was the most convenient in counting micro-colonies. Phasecontrast micrographs not only eliminate the use of stain to distinguish between live and dead cells but also help us identify contaminants to some extent. The advantages to this technique are many, but it too has some cons. Phase-contrast microscope itself comes with a hefty price tag but this method of CFU counting can be performed using any microscope that comes with a scale to extrapolate. Also, this technique demands manual labour; albeit lesser labour-intensive than the traditional counting ways. It is pertinent to note that our method is in support of the method published earlier by Thomas et al (2015) in which they counted bacterial and yeast CFU using single plate serial dilution spotting. However, in their study, no micro-colony approaches were made.

Microliter spotting and micro-colony observation were used to calculate generation time in *E. coli*. Generation time was observed to be 25 min. This further suggested the useful nature of this method. We further used it to study the antibiotic-resistant mutation rate in *E. coli*. By subsequent plating of *E. coli* culture in LB-Agar supplemented with rifampicin and counting mutant colonies at every 1 h interval, we obtained the mutation rate at the intervals of 2 h – 3 h, 2 h – 4 h and 3 h – 4 h as 1.73 × 10^-8^, 1.09 × 10^-8^, 2.27 × 10^-8^ respectively [Table 6]. Interestingly, while performing the mutation rate study for rifampicin resistance in *E. coli*, sometimes a negative mutation rate was observed, which might be due to (i) having slower growth in mutant colonies compared to wild-type colonies. (ii) competition among mutant strains and the wild-type strain. (iii) larger generation time than the wild-type. An elaborate study is needed to further analyse these findings on the mutation rate.

**Table 5.** Mutation frequency.

| Time | Bacterial populations (CFU/ml) | Avg. colonies on Rif<sup>r</sup> plates | Rif<sup>r</sup> frequency |
| --- | --- | --- | --- |
| 1 h | 1.44 x 10<sup>8</sup> | 6.6 | 4.58 x 10<sup>-8</sup> |
| 2 h | 7.33 x 10<sup>8</sup> | 16.66 | 2.28 x 10<sup>-8</sup> |
| 3 h | 6.66 x 10<sup>9</sup> | 250 | 3.78 x 10<sup>-8</sup> |
| 4 h | 2.22 x 10<sup>10</sup> | 1700 | 7.65 x 10<sup>-8</sup> |

**Table 6.** Mutation rate.

| Time | Mutation rate in Rif<sup>r</sup> *E. coli* |
| --- | --- |
| 2 h - 3 h | 1.73 x 10<sup>-8</sup> |
| 2 h - 4 h | 1.09 x10<sup>-8</sup> |
| 3 h - 4 h | 2.27 x10<sup>-8</sup> |

The fluctuation test was performed in analysing Rif^r^ mutants in *E. coli*. The number of Rif^r^ mutants in different samples from the same culture and different samples from a series of similar cultures were analysed [Table 7; Fig. 6]. The variance was 21.94 in the case of different samples from a series of similar cultures whereas the same was 6.57 in the case of different samples from the same culture. The higher variance in the case of different samples from a series of similar cultures was in concordance with the observation made earlier by Luria and Delbruck in their fluctuation test that explained that genetic mutations arise in the absence of selective pressure rather than being a response to it. We believe that the simple approach that has been demonstrated in enumerating bacterial CFU is going to be helpful to researchers.

**Table 7.**
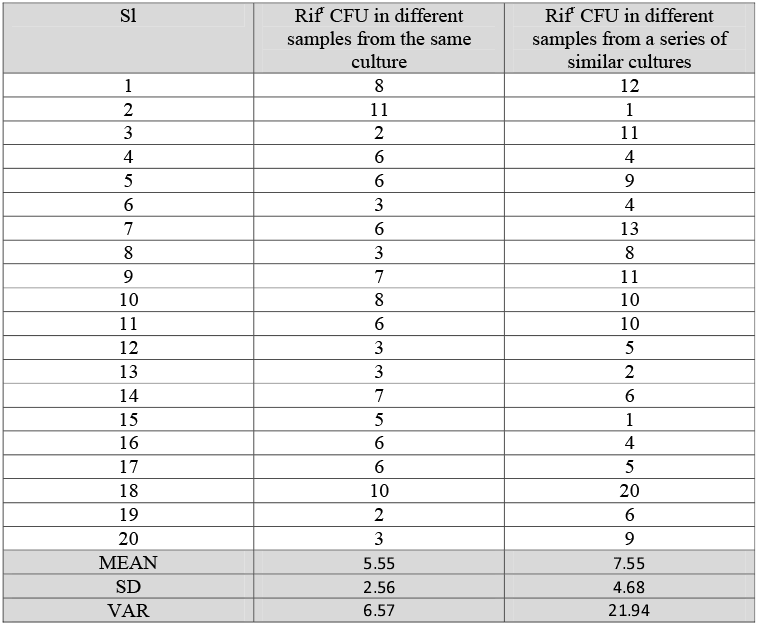
Fluctuation test.

**Fig 5.**
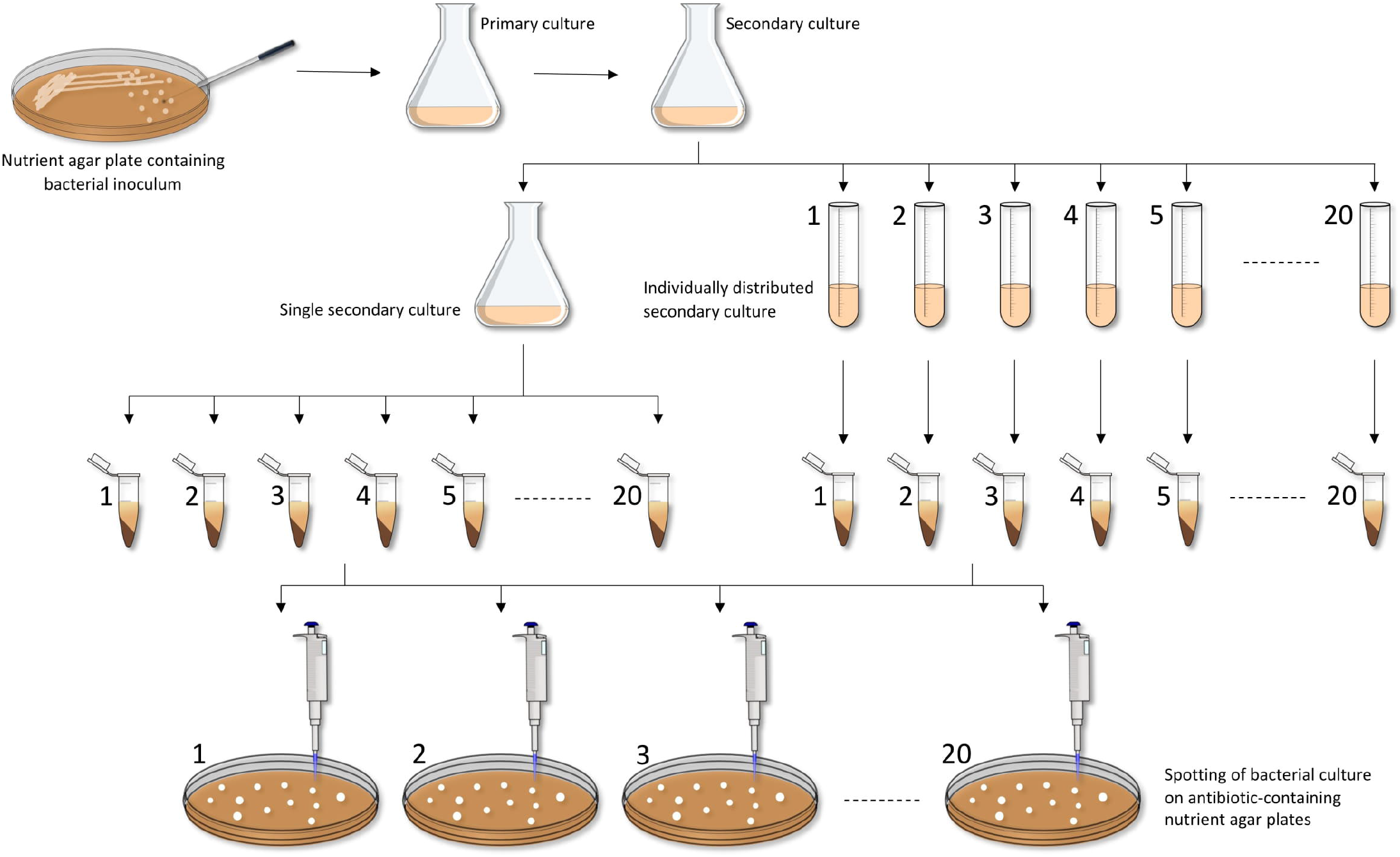
Fluctuation test procedure.

**Fig 6.**
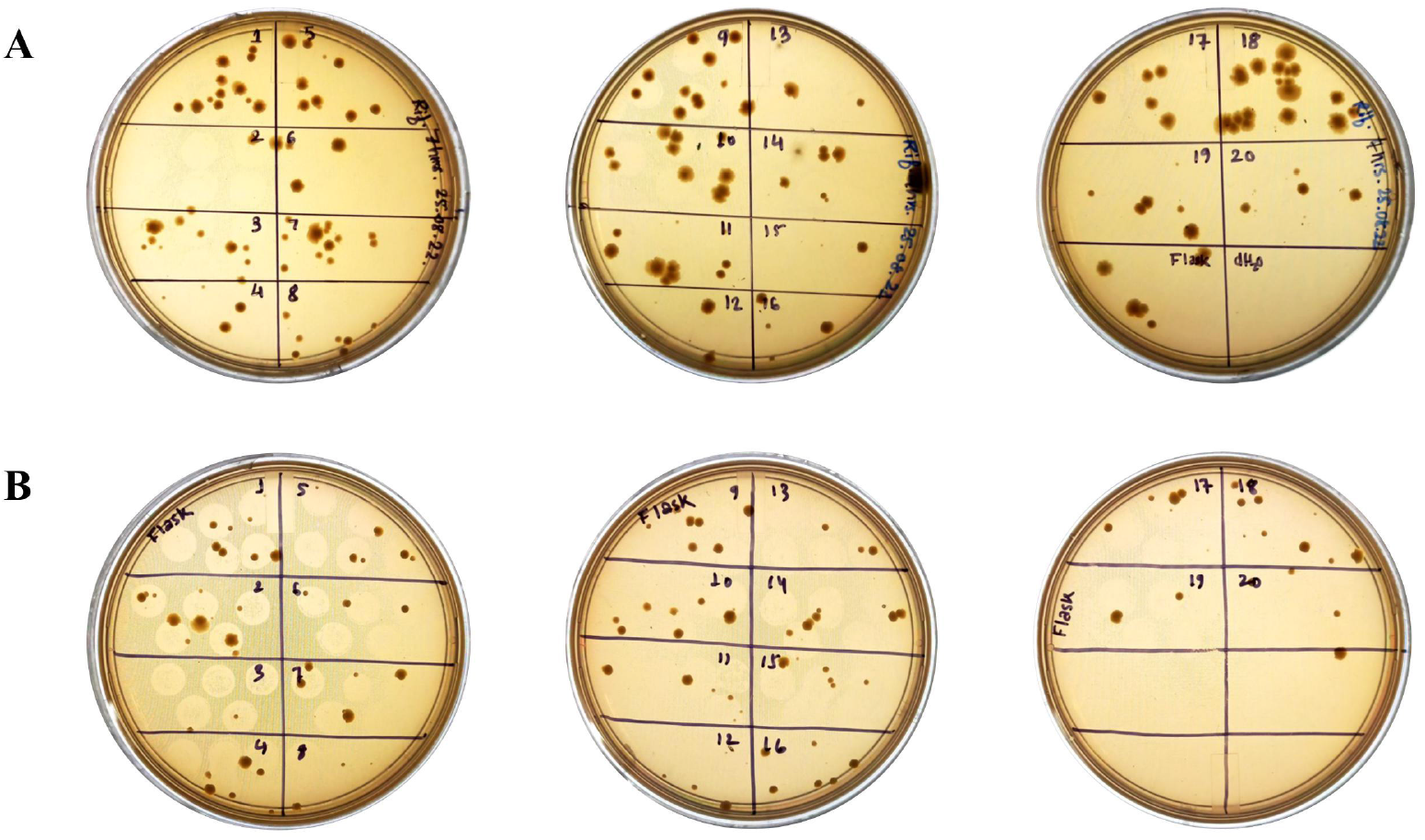
Fluctuation test in *E. coli* by the micro-colony counting method. A. Colony count of mutants obtained from different colonies of the same culture, B. Colony count of mutants obtained from samples of a series of similar cultures.

## Acknowledgements

SB is thankful for the JRF/SRF fellowship from the UGC-NFSC, GoI, New Delhi. MY is thankful for RA fellowship from the SERB-DST grant (CRG/2020/002651) awarded to AK. SS is thankful to CSIR, GoI, New Delhi for the JRF/SRF fellowship. SJG is thankful for the JRF fellowship from the DBT, GoI New Delhi grant (BT/PR41637/NER/95/1753/2021) awarded to SKR, and also to Tezpur University for the Institutional fellowship. S Begum is thankful to Tezpur University for the institutional fellowship. AJ is thankful to UGC, GoI, New Delhi for the NET-JRF fellowship. SD is thankful to DBT, GoI, New Delhi for the fellowship during his master’s programme.

## Funding

The authors would like to acknowledge DBT for the MSc project grants for the consumables and DBT-U-Excel/NER for the equipment.

## Conflict of interest

We declare that there is no conflict of interest among the authors of this manuscript.

## Notes

### Competing Interest Statement

The authors have declared no competing interest.

### Summary of Updates

Typos removed

