## Supplementary Figures for "Microliter spotting and micro-colony observation: a rapid and simple approach for counting bacterial colony forming units"

SFig 1(a). Micro-colonies of *E. coli* cells (10^4^ dilutions) on LB agar plates were observed after **(A)** 2 h **(B)** 4 h **(C)** 6 h **(D)** 8 h incubation at 37 ᴼC.

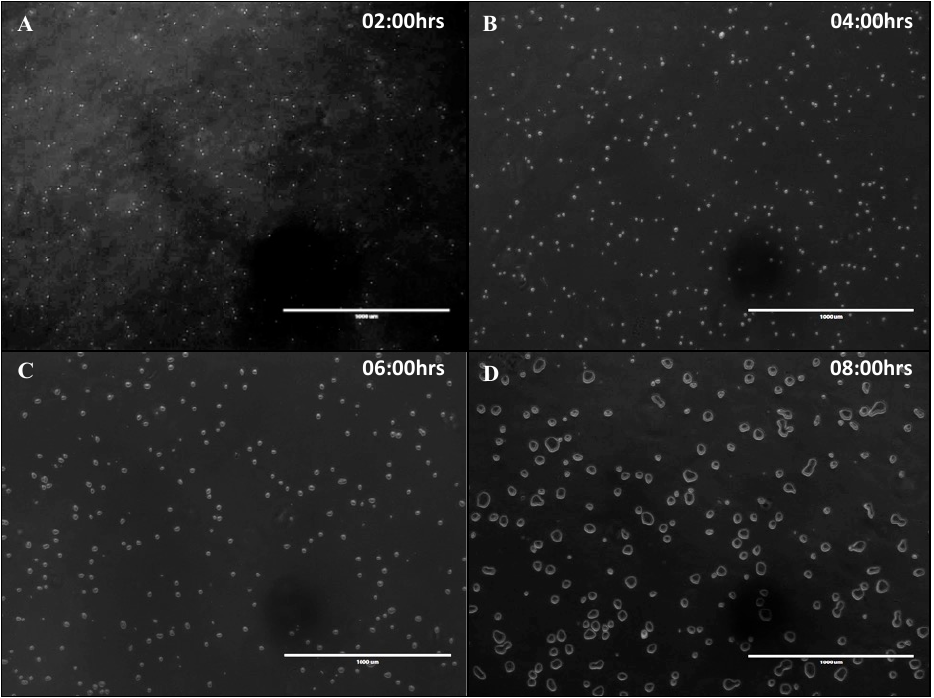

SFig 1(b). Micro-colonies of *E. coli* cells (10^5^ dilutions) on LB agar plates were observed after **(A)** 2 h **(B)** 4 h **(C)** 6 h **(D)** 8 h incubation at 37 ᴼC.

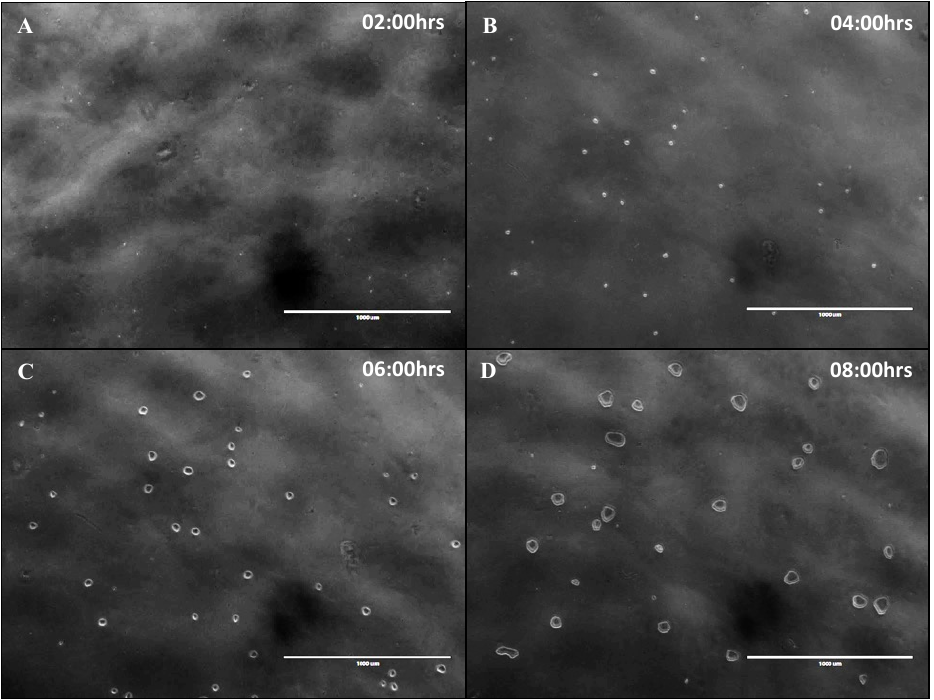

SFig 1(c). Micro-colonies of *E. coli* cells (10^6^ dilutions) on LB agar plates were observed after **(A)** 4 h **(B)** 6 h **(C)** 8 h at 37 ᴼC.

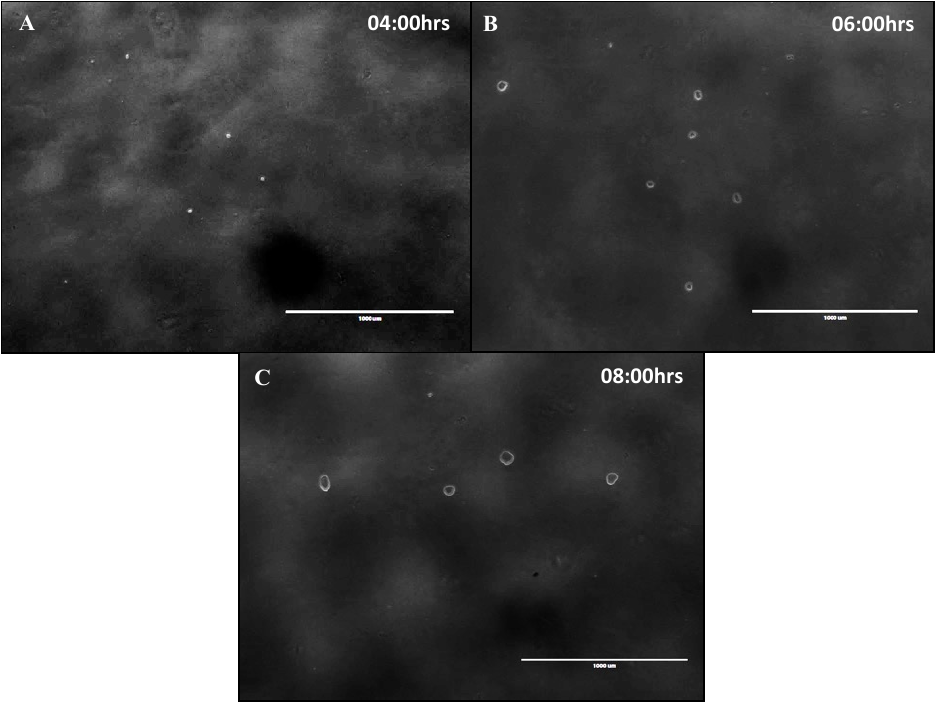

SFig 2(a). Micro-colonies of *R. solanacearum* cells (10^4^ dilutions) on BG agar plates were observed after **(A)** 8 h **(B)** 10 h **(C)** 12 h **(D)** 14 h incubation at 28 ᴼC.

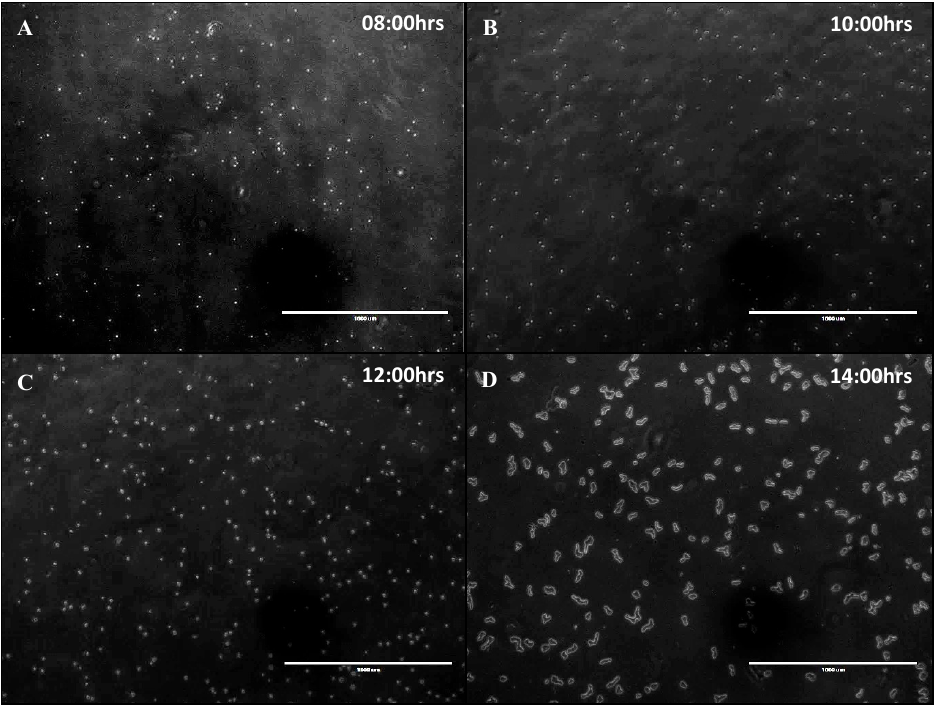

SFig 2(b). Micro-colonies of *R. solanacearum* cells (10^5^ dilutions) on BG agar plates were observed after **(A)** 8 h **(B)** 10 h **(C)** 12 h **(D)** 14 h incubation at 28 ᴼC.

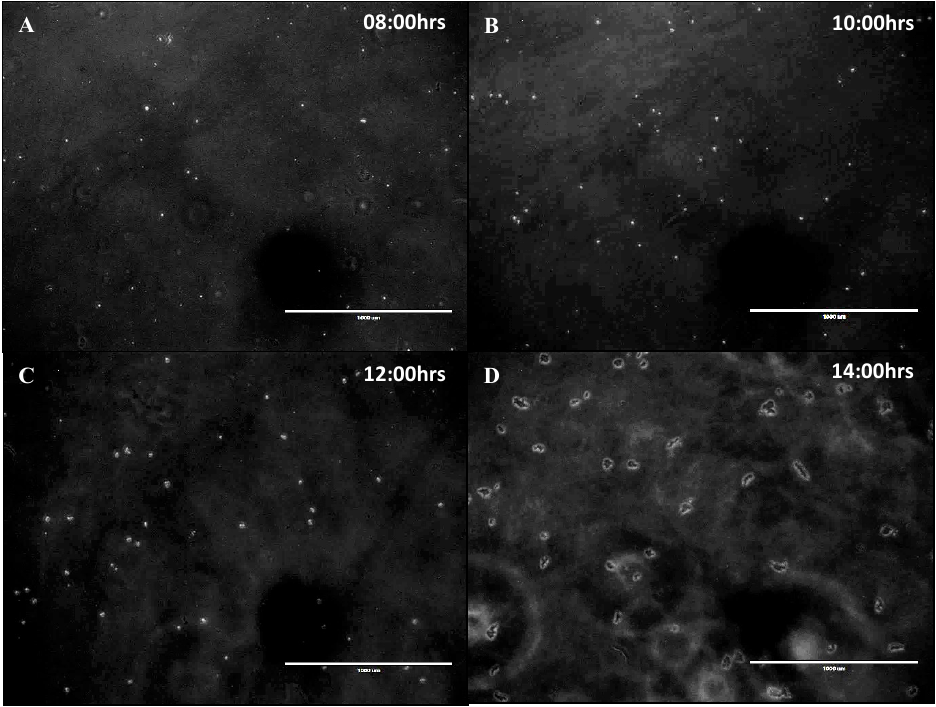

SFig 2(c). Micro-colonies of *R. solanacearum* cells (10^6^ dilutions) on BG agar plates were observed after **(A)** 8 h **(B)** 10 h **(C)** 12 h **(D)** 14 h incubation at 28 ᴼC.

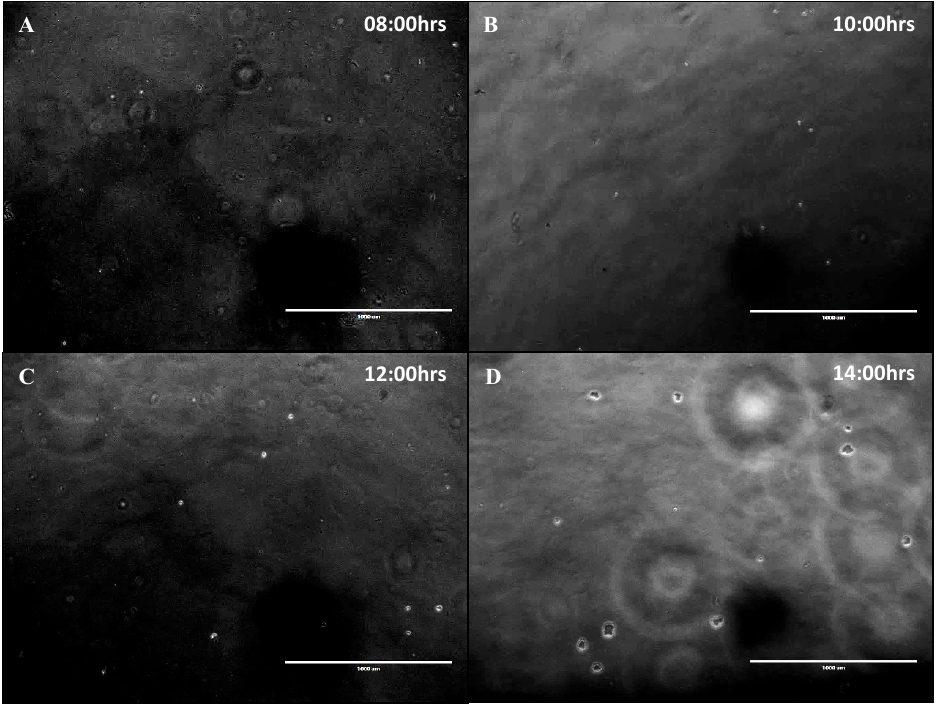

SFig 3. By spread plate method, visible colonies of *R. solanacearum* F1C1(A to D) and *E. coli* DH5α cells (E to H) were observed in various dilutions.

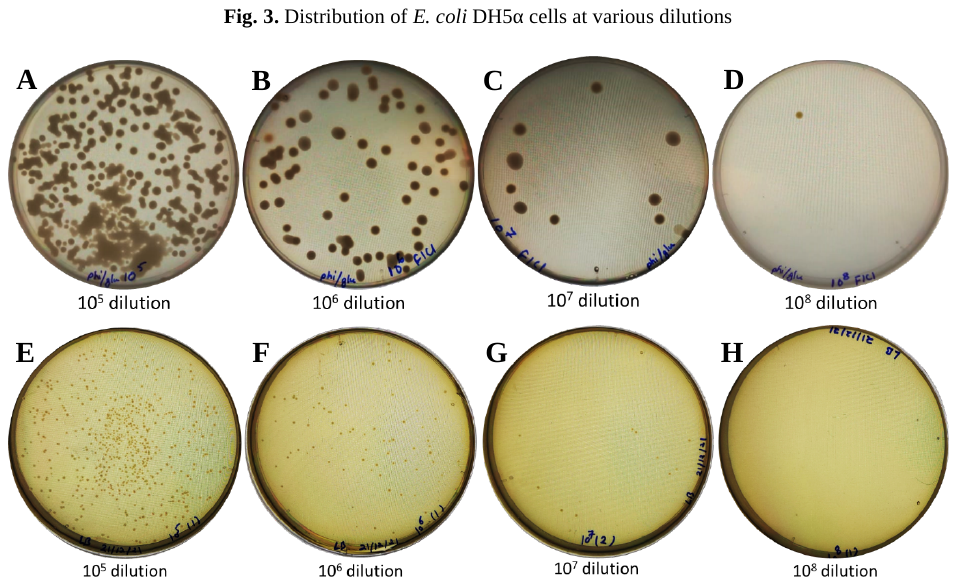

SFig 4. *R. solanacearum* CFU counting by simple dilution method

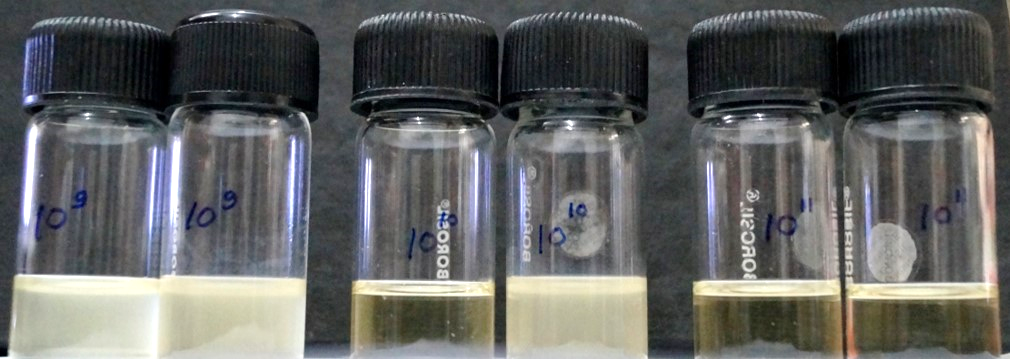

A saturated culture of *R. solanacearum* was diluted as 10^1^, 10^2^, 10^3^ ----- 10^9^, 10^10^, 10^11^, 10^12^ folds in the growth medium. Three culture tubes in each dilution were maintained. Turbidity appeared in the medium and was observed for five days. In 10^8^ dilutions turbidity appeared by 48 h in each tube. In 10^9^ dilutions turbidity appeared by 72 h in each of the three tubes, but in 10^10^ dilutions only in one tube out of the three turbidity was observed by 72 h. Till 5 days period, no turbidity was observed in case of 10^11^ and 10^12^ dilutions as well as two of the three tubes in case of 10^10^ dilution. So, without plating, it can be said that a minimum of 10 CFU are present in the culture.

SFig 5. By spread plate method, visible colonies of rifampicin resistance *E. coli* BL21 (A to D) and spectinomycin resistance (E to H) were observed from 1 h to 4 h.

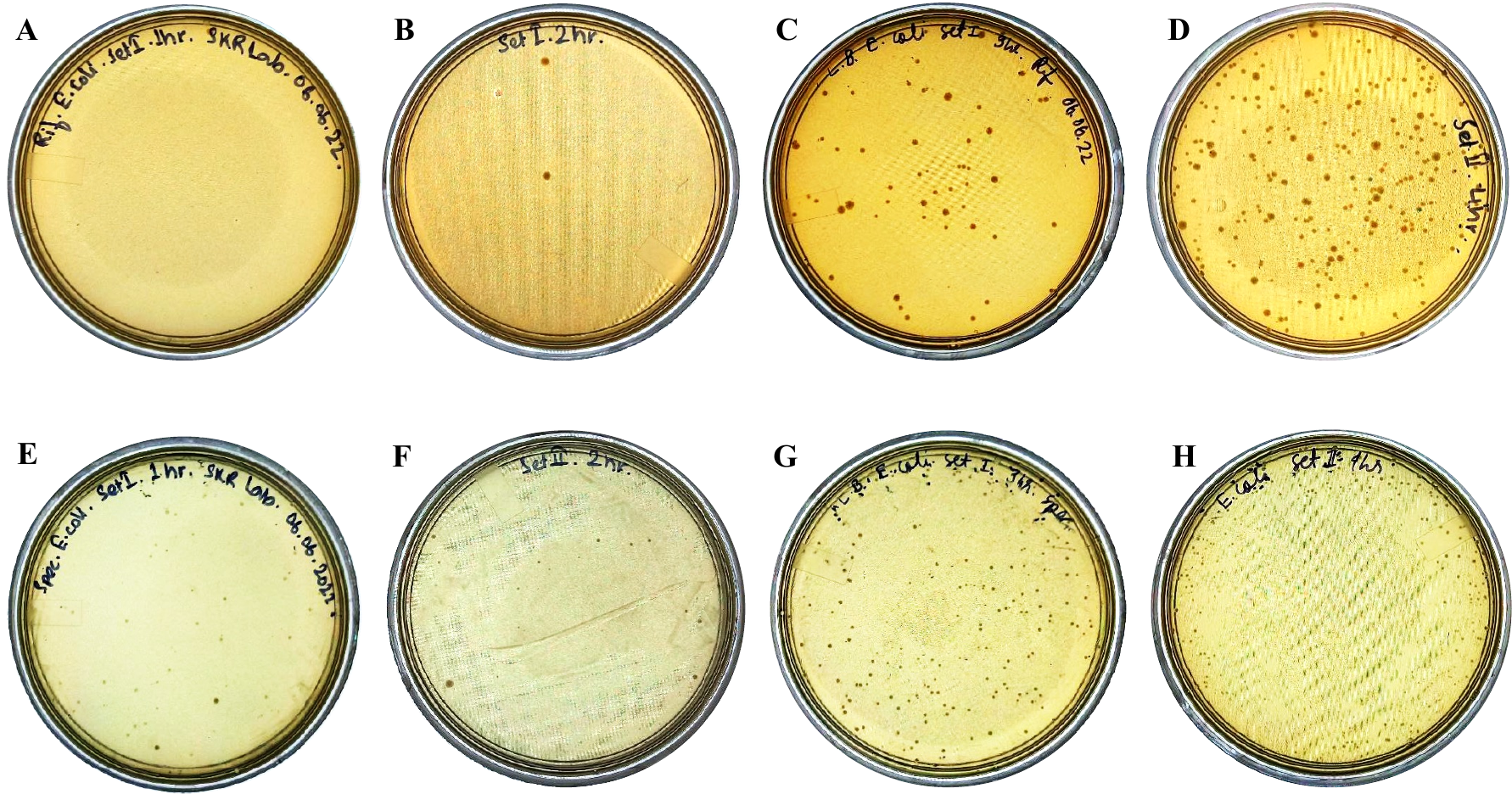
